# Crystal violet structural analogues identified by *in silico* drug repositioning present anti-*Trypanosoma cruzi* activity through inhibition of proline transporter TcAAAP069

**DOI:** 10.1101/645333

**Authors:** Melisa Sayé, Lucrecia Gauna, Edward Valera-Vera, Chantal Reigada, Mariana R. Miranda, Claudio A. Pereira

## Abstract

**Background:** Crystal violet (CV) was used for several years in blood banks to eliminate the parasite *Trypanosoma cruzi* in endemic areas in order to prevent transfusion-transmitted Chagas disease. One mechanism of action described for CV involves inhibition of proline uptake. In *T. cruzi*, proline is essential for host cell infection and intracellular differentiation among other processes, and can be obtained through the proline permease TcAAAP069.

**Methodology/Principal Findings:** CV inhibited proline transporter TcAAAP069 and parasites overexpressing this permease were 47-fold more sensitive to this compound than control parasites. Using CV as reference molecule, loratadine, cyproheptadine, olanzapine and clofazimine were identified as structurally related compounds to CV (structural analogues) by *in silico* drug repurposing through a similarity-based virtual screening protocol. All these already-approved drugs for clinical use inhibited TcAAAP069 activity with different efficacies and also presented trypanocidal action in epimastigotes, trypomastigotes and amastigotes of the Y strain. Additionally, loratadine, cyproheptadine and clofazimine showed trypanocidal effect on epimastigotes of the CL Brener and DM28c strains. Finally, a synergistic effect between benznidazole and the CV chemical analogues was evidenced by combination and dose-reduction indexes values in epimastigotes of the Y strain.

**Conclusions/Significance:** Loratadine, cyproheptadine and clofazimine inhibit TcAAAP069 proline transporter and also present trypanocidal effect against all *T. cruzi* life stages. These CV structural analogues could be a starting point to design therapeutic alternatives to treat Chagas disease by finding new indications for old drugs. This approach, called drug repurposing is a recommended strategy by the World Health Organization to treat neglected diseases, like Chagas disease, and combination therapy may improve the possibility of success of repositioned drugs.

**Author summary:** Chagas disease, caused by the parasite *Trypanosoma cruzi*, affects 7 million people worldwide. Despite there are two drugs available since 50 years ago, the therapy present severe side effects and is not effective in the chronic phase of the disease were most of the patients are diagnosed. Crystal violet (CV) was utilized as additive in blood banks to prevent transfusion-transmitted Chagas disease. Proline is involved in many pathways, like infection establishment and life cycle progression. In this work we first demonstrate that CV has the proline permease TcAAAP069 as one of its molecular targets. Then we search in a database of already-approved drugs for compounds that were structurally related to CV under the premise “similar structure, similar activity”. We identified three drugs that inhibit proline transport and present at least the same trypanocidal effect than benznidazole, the current treatment for Chagas disease. Finally we observed a synergistic effect with the multidrug combination therapy. Drug discovery is an expensive and time-consuming process and Chagas disease is associated with poverty. The discovery of new indications to old drugs, called drug repurposing, can facilitate a rapid and more profitable therapy application since preclinical trials and pharmacokinetic studies are already available.

## Introduction

Chagas disease is a neglected disease caused by the protozoan parasite *Trypanosoma cruzi* that can mainly be acquired through an insect vector, blood transfusion and placental or congenital transmission. Among these mechanisms, blood transfusion is still one of the most important mechanisms of transmission of Chagas disease due to lack of blood bank control and migration of people from endemic to non-endemic countries (1,2). Crystal violet (CV) is a well-known dye common in microbiology and it was used for a long time as an additive in blood banks as an attempt to eliminate Chagas transmission in endemic areas (3). Different trypanocidal mechanisms have been proposed for CV, among them Hoffmann *et al.* demonstrated that proline and methionine transport were strongly inhibited by this compound (4).

Chagas disease affects 7 million people in Latin America and almost 70 million more are at risk of infection (5). The only two approved drugs to treat this disease, benznidazole and nifurtimox, were discovered over 50 years ago and present limited efficacy and several side effects (6). Thus the development of new therapies to treat Chagas disease remains as an urgent need.

The first multigenic family of amino acid transporters from *T. cruzi* (TcAAAP; Amino Acid/Auxin Permeases; TC 2.A.18) was identified by our group (7). Few members of this family have been characterized in trypanosomatids, including the proline (8) (TcAAAP069), arginine (9,10) (TcAAP3) and lysine (11) (TcAAP7) permeases. A polyamine transporter (12,13) (TcPAT12) has also been characterized in *T. cruzi* and it probably belongs to the Amino Acid-Polyamine Organocation (APC; TC 2.A.3) family, which is phylogenetically very close to the AAAP family. The transporter TcAAAP069 is mono-specific for D- and L-proline and it is involved in resistance mechanisms to trypanocidal drugs and also to reactive oxygen species including hydrogen peroxide and nitric oxide (8). In addition, proline is a relevant amino acid for trypanosomatids since it is used as an alternative carbon and energy source to glucose (14). It has been proved that in *T. cruzi* proline confers osmotic stress resistance (15) and it also sustains cellular invasion in metacyclic trypomastigotes (16) as well as differentiation of intracellular epimastigotes to trypomastigote forms, which is required for the establishment of infection in mammalian hosts (17).

The orthologous genes of TcAAAP069 have been characterized in other trypanosomatids. In *T. brucei*, the orthologous permease TbAAT6 is a low-affinity low-selective transporter for neutral amino acids, including proline (18). The loss of TbAAT6 is sufficient to develop resistance to eflornithine, a drug used to treat human African trypanosomiasis (HAT) (19). Moreover, the importance of amino acid transporters in *T. brucei* has been demonstrated by RNAi-mediated knock-down of arginine and lysine transporters TbAAT5-3 and TbAAT16-1 from the AAAP family which resulted in growth arrest in bloodstream forms (20). In *Leishmania donovani*, the transporter LdAAP24 mediates proline and alanine uptake and regulates the response to osmotic stress and amino acid homeostasis (21). The parasite *L. donovani* naturally expresses two variants of the proline/alanine transporter, one 18 amino acids shorter than the other. While in *T. cruzi* no information is available about the determinants of the substrate specificity of TcAAAP transporters, in LdAAP24 these 18 amino acids define its substrate specificity for both amino acids (LdAAP24 long variant) or for proline alone (Δ18LdAAP24 short variant) (22). Recently, it has been reported that LdAAP24 also presents a stage-specific expression and it is rapidly degraded when promastigotes are differentiated to amastigotes (23). TcAAAP069 activity and localization have been studied only in epimastigotes overexpressing this proline permease (24), thus there is no information regarding stage-specific expression.

Since the transporter TcAAAP069 has many differential features compared to those present in mammals and proline is essential for parasite survival, it could be possible to design a specific inhibitor of this *T. cruzi* permease with trypanocidal action and non-toxic for the host cells. In fact, a synthetic proline analogue named ITP-1G has been validated as a TcAAAP069 inhibitor which also possesses trypanocidal effect (25). One of the best strategies for the identification of new treatments for neglected diseases is drug repositioning, which is the process of finding new indications for existing drugs. The main benefit of this approach relays on those drugs being ahead in the development and clinical phase I trials, consequently they could reach the patients in approximately half of the time and with a significant lower cost (26). Some successful drugs repositioned for neglected diseases are eflornithine and miltefosine which were originally discovered for cancer therapy and now are used to treat HAT and leishmaniasis, respectively (27). Considering this information, in this work we performed an *in silico* screening of a drug database to find compounds structurally related to CV that could also inhibit TcAAAP069 and exert trypanocidal effect with less toxicity.

## Methods

### Parasites

Epimastigotes of Y, CL Brener and Dm28c strains (5 x10^6^ cells/mL) were cultured at 28 °C in plastic flasks (25 cm^2^), containing 5 mL of BHT (brain-heart infusion-tryptose) medium supplemented with 10% fetal calf serum (FCS), 100 U/mL penicillin, 100 μg/mL streptomycin and 20 μg/mL hemin (28). Trypomastigotes were obtained from the extracellular medium of VERO infected cells as previously described (17).

### Cells

VERO cells (African green monkey Kidney, ATCC® CCL-81™) were cultured in MEM medium supplemented with 10% heat inactivated FCS, 0.15% (w/v) NaHCO_3_, 100 U/mL penicillin and 100 U/mL streptomycin at 37°C in 5% CO_2_.

### TcAAAP069 overexpressing parasites

The TcAAAP069 overexpressing parasites were available in our laboratory and were previously characterized (25). Briefly, TcAAAP069 (TriTrypDB ID: TcCLB.504069.120) was amplified using genomic *T. cruzi* DNA as template. Amplification product was subcloned into a modified pTREX expression plasmid called pTREXL (pTREX-069) (29,30). Constructions were transfected into *T. cruzi* epimastigotes of the Y strain as previously described (31).

### Transport assays

Aliquots of epimastigote cultures (10^7^ parasites) were centrifuged at 8,000 xg for 30 s, and washed once with phosphate-buffered saline (PBS). Cells were resuspended in 0.1 mL PBS and the assay started by the addition of 0.1 mL of the transport mixture containing [^3^H]-proline (PerkinElmer’s NEN^®^ Radiochemicals; 0.4 μCi) in the presence of different concentrations of the indicated drug. Following incubation at 28 °C, reaction was stopped by adding 1 mL of ice-cold PBS. Cells were centrifuged as indicated above, and washed twice with ice-cold PBS. Cell pellets were resuspended in 0.2 mL of water and counted for radioactivity in UltimaGold XR liquid scintillation cocktail (Packard Instrument Co., Meridien CT, USA) (32,33). Cell viability was assessed by direct microscopic examination. Non-specific uptake and carry over were assayed without incubation (T_0_), or incubated at 4 °C.

### Trypanocidal activity assays

Epimastigotes of *T. cruzi* were cultured as described above, in 24-wells plate at a start density of 0.5 x10^7^ cells/mL in BHT medium. Parasites′ growth was evaluated at different concentrations of BZL, CV and CV structural analogues, and parasite proliferation was determined after 48 h. Trypanocidal activity in trypomastigotes and amastigotes was performed using 1 x10^6^ cells/mL in 96-well plates and incubating at 37 ̊C for 24 h in the presence of the corresponding drug. Growth was determined by counting in a Neubauer chamber or by viability assays using “Cell Titer 96^®^ Aqueous One Solution Cell Proliferation Assay (MTS)” (Promega, Madison, WI, USA) according to the manufacturer’s instructions.

### Toxicity assays in VERO cells

The toxicity of the compounds was determined in VERO cells by the CV staining assay. The cells (1 x10^4^ cells/well) were incubated in 96-well plates with the indicated compound or DMSO as negative control and maintained at 37° C for 24 h. Then the cells were fixed for 15 min, and stained with 0.5% CV. The absorbance was measured at 570 nm. The selectivity index (SI) was calculated for each CV analogue as the ratio between the IC_50_ obtained in mammalian cells and the IC_50_ in trypomastigotes. For the combination therapy, the results were presented as percentage of untreated cells.

### Virtual screening

Similarity screening search was performed using the CV structure as reference (query) compound (ZINC ID: 13763987). Structures (≈ 10,000) were obtained from the highly curated “Sweetlead” database of world’s approved medicines, illegal drugs, and isolates from traditional medicinal herbs (34). The screening was performed using the softwares LiSiCA v1.0 (35) (Ligand Similarity using Clique Algorithm), KCOMBU (36) (K(ch)emical structure COMparison using the BUild-up algorithm), ShaEP (37) (Molecular Overlay Based on Shape and Electrostatic Potential) v 1.1.2, and Shape-it v 1.0.1 (38) (Gaussian volumes as descriptor for molecular shape). For all of them different control structures were used in screening searches in order to check they appear with the higher score in each resulting list, validating the retrieve capabilities of the programs. Then, the screening with CV was performed.

### Docking simulations and structural alignments

The homology model of TcAAAP069 was constructed using the I-TASSER-server (39) and was based mainly on the structure of the *Escherichia coli* arginine/agmatine antiporter AdiC (PDB ID: 3LRB), in addition to 9 templates of other proteins showing coverage between 0.72 and 0.95. The modeled structure had 93.7% residues in the residues in favored region of Ramachandran’s plot, and none of the residues located in non-allowed regions were involved in the active site of the transporter. The model was refined using the software Modeller v7 (40). AutoDock 4.2.6 (41) was employed for docking assays, using a grid that covers the whole transporter molecule to calculate the optimal energy conformations for the ligands interacting with the residues within the permease cavity. The program was run using a Lamarckian Genetic Algorithm 100 times, with a population size of 300, and 2.7 x10^4^ as maximum number of generations. Structures alignments were performed using LigandScout 4.1.10 (42) (Inte:Ligand, Software-Entwicklungs und Consulting GmbH, Austria) whose license was kindly granted to our laboratory by the company. Modeling was performed using the LigandScout default parameters.

### Analysis of synergism

Calculations were performed using CompuSyn software (http://combosyn.com) which is based on the median-effect equation (43) and its extension, the combination index (44). Results are reported as Combination Index (CI), where CI < 1, CI = 1 and CI > 1 indicate synergism, additive effect and antagonism, respectively. LTD, CPH and CFZ were evaluated as a single drug (LTD-CPH-CFZ) in combinations according to the following scheme: 1/2 IC_50_, 1/3 IC_50,_ 1/4 IC_50_,> 1/5 IC_50_, 1/10 IC_50_ and 1/25 IC_50_, where IC_50_ refers to the value obtained for each one of the three CV analogues (25 μM, 50 μM and 10 μM of LTD, CPH and CFZ, respectively for epimastigotes). 1/2 IC_50_ means 42.5 μM (the sum of half of the IC_50_ value for each CV analogue, 12.5 μM + 25 μM + 5 μM, for LTD, CPH and CFZ, respectively), 1/5 IC_50_ means 17 μM (5 μM + 10 μM + 2 μM), 1/10 IC_50_ means 8.5 μM (2.5 μM + 5 μM + 1 μM) and 1/25 IC_50_ means 3.4 μM (1 μM + 2 μM + 0.4 μM). BZL was evaluated at the following concentrations: 10 μM, 5 μM, 1 μM and 0.1 μM.

### Statistics and data analysis

IC_50_ values were obtained by non-linear regression of dose-response logistic functions, using GraphPad Prism 6.01 for Windows, and synergism was evaluated using the free software CompuSyn. All experiments were performed in triplicate and the results are presented as mean ± standard deviation (SD).

## Results

### Effect of CV on the proline transporter TcAAAP069

In order to test if the previously reported decrease in proline transport by CV (4) is due to the specific inhibition of the TcAAAP069 proline permease, the effect of CV on this transporter activity was evaluated. After incubation with CV, doses that inhibit 50% of the proline transport (IC_50_) were calculated for wild type (TcWT) and TcAAAP069 overexpressing (Tc069) *T. cruzi* epimastigotes. The inhibitory effect of CV was significantly decreased in Tc069 parasites with a calculated IC_50_ 2.5-fold higher than control parasites (7.1 μM ± 0.8 and 17.7 μM ± 1.1, respectively; p<0.001) (Fig 1a).

**Fig 1.**
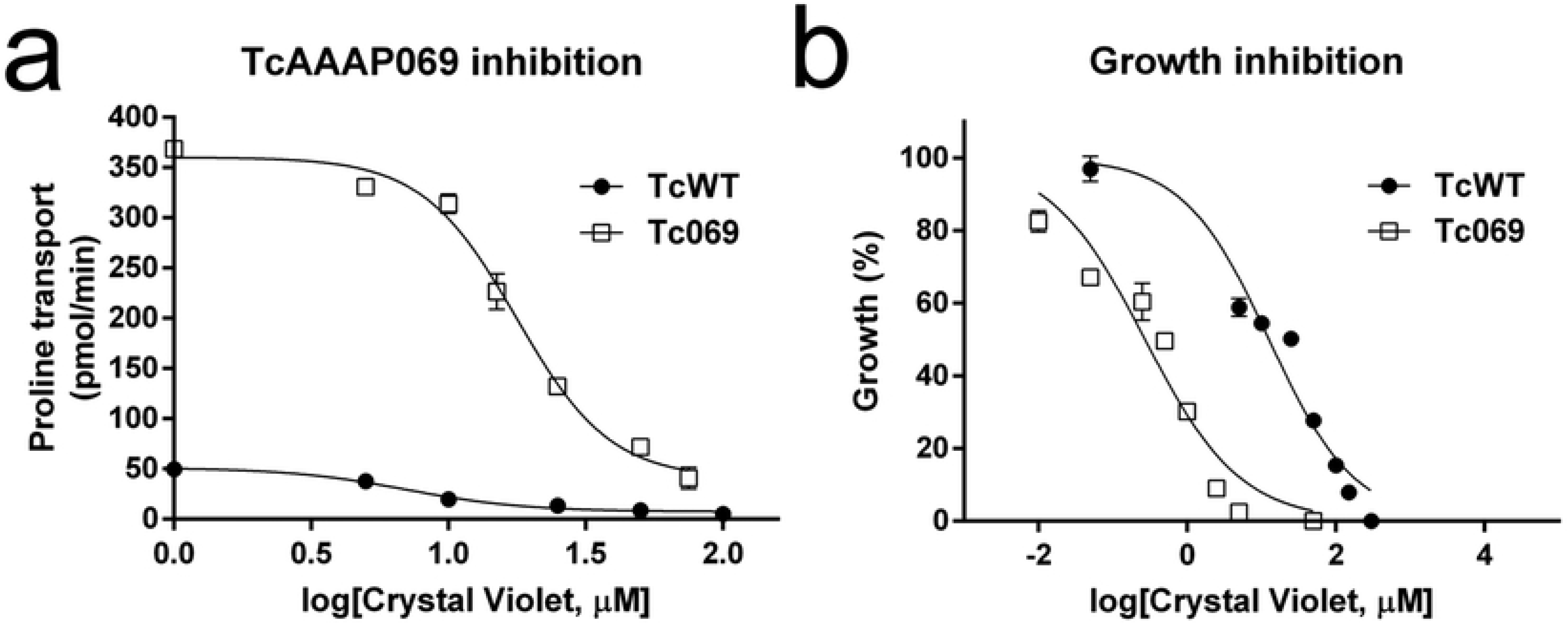
Validation of crystal violet as proline transporter TcAAAP069 inhibitor. Inhibition of proline transport in wild type parasites (TcWT, •) and parasites overexpressing TcAAAP069 (Tc069, □). (b) Trypanocidal effect of crystal violet in TcWT (•) and Tc069 (□) parasites. The data is expressed as the mean ± standard deviation and corresponds to one of three independent experiments.

To further demonstrate that CV specifically interacts with TcAAAP069, the trypanocidal effect of CV was evaluated using the same parasite models (TcWT and Tc069 cultures). The toxicity of CV significantly increased when TcAAAP069 was overexpressed as evidenced by a reduction in the IC_50_ value of 47-fold compared to wild type controls (0.27 μM ± 0.06 and 12.67 μM ± 2.1, respectively; p<0.0001) (Fig 1b). All these evidences demonstrate that proline transporter TcAAAP069 is one of the targets of CV.

### Similarity virtual screening

In order to identify chemical analogues of CV approved for human administration, the Sweetlead database was analyzed using four different algorithms that rank molecules according to their similarity (LiSICA, ShaEP, KCOMBU and Shape-it). Compounds that complied with the following criteria were selected: 1) being in the top five scored according to at least two different algorithms; 2) have been used in humans, 3) sharing a substructure with CV by 3D alignment, and 4) being easily accessible in the pharmaceutical companies. Only four compounds fitted all the established conditions; these drugs were loratadine (LTD; antihistamine), olanzapine (OLZ; antipsychotic), cyproheptadine (CPH; antihistamine) and clofazimine (CFZ; antibiotic). Dapsone (DPS; antibiotic) is an already-approved drug that shares some substructures with CV, but since it was not in the top scored results it was designated as a control of the selection strategy. The chemical structures of CV and its analogues are presented in the Fig 2. It is worth mentioning that the result with the highest score obtained with the four programs used, was the reference molecule (CV). To identify the common chemical features between CV and the selected chemical analogues, the LigandScout algorithm was used in order to perform feature-based structure alignments where the similarities are calculated as the number of matched feature pairs (MFP; i.e. aromatic ring, hydrophobic area, hydrogen bond donor or acceptor, negative or positive ionizable atom and metal binding location). The CV possesses eight features that were set as references, which are four hydrophobic interactions, three aromatic rings and one positive ionizable interaction. The chemical features shared between CV and its chemical analogues as well as the alignments are schematized in the S1 Fig. The four CV analogues, CPH, LTD, CFZ and OLZ, possess between 3 and 5 MFPs out of a total of 8 present in the CV. CPH is the compound that shared more chemical features with CV; including two hydrophobic interactions, two aromatic rings and one positive ionizable interaction.

**Fig 2.**
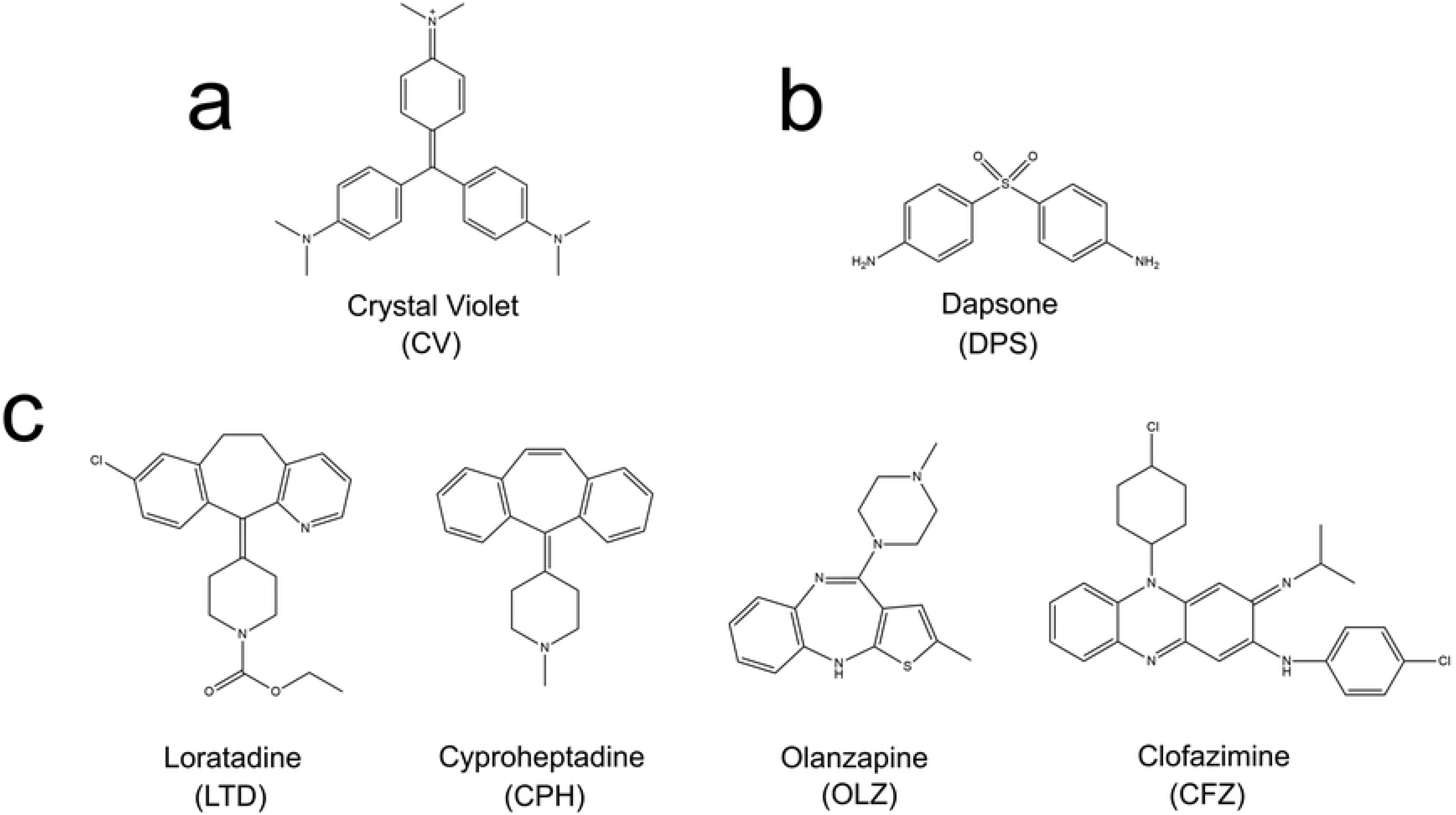
Chemical structures of crystal violet, its analogues and negative control. The figure shows the chemical structure of crystal violet (a), the structure of the negative control, dapsone, (b) and the structures of the four crystal violet structural analogues loratadine, cyproheptadine, olanzapine and clofazimine (c).

### Molecular docking of the TcAAAP069 inhibitors

The possible modes of interaction between LTD, CPH, OLZ, and CFZ with the proline transporter TcAAAP069 were tested by a simulation with AutoDock 4.2.6, using proline and CV as a reference of experimentally identified high affinity binders and to determine if the structure similarity of the selected compounds and the CV translates into a similar mode of interaction with the protein. To check if, as expected, there were not predicted interactions between the transporter and a molecule that does not comply with the selection criteria, a simulation with DPS was also performed. According to the obtained docking models, proline, CV and the four analogues bind different residues groups, located within the permease channel while DPS does not appear to bind to any pocket in TcAAAP069 (Fig 3a). The lowest score values (ΔG) of the predicted interaction of proline and CV with the transporter are −4.07 and −5.26 kcal.mol^−1^, respectively. The binding scores of the four CV structural analogues were lower than for proline and CV, with values between −9.60 kcal.mol^−1^ to −5.81 kcal.mol^−1^. This suggests that CV and its chemical analogues form a more stable bond to that channel region of TcAAAP069 than the natural ligand. All these data are summarized in Table 1.

**Fig 3.**
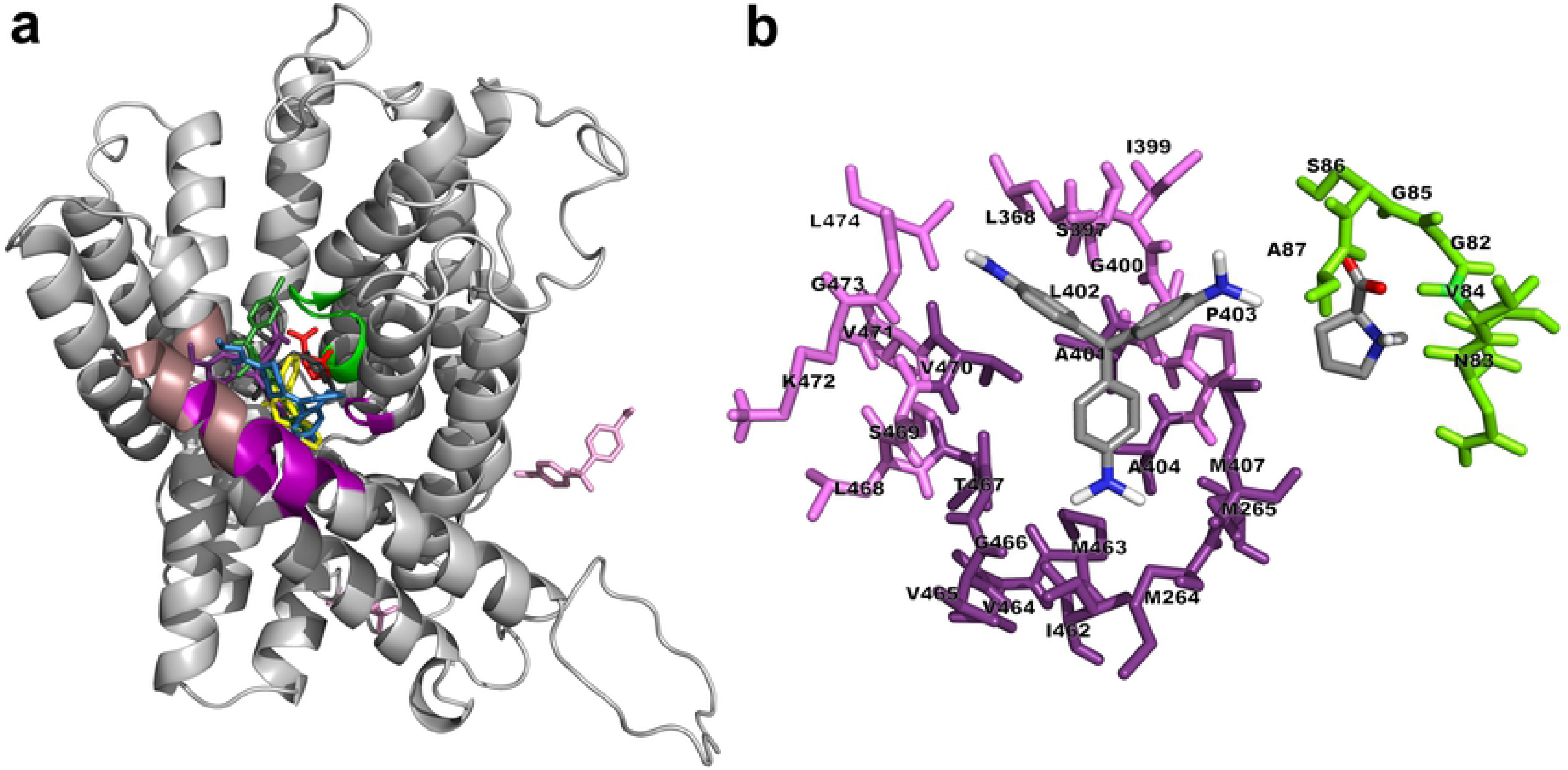
Binding poses suggested by molecular docking between proline, crystal violet and its chemical analogues with proline permease TcAAAP069. (a)Homology modelled TcAAAP069 (represented as cartoon) highlighting the predicted interaction sites of proline (PRO site, green) and crystal violet (CV site, violet; and dark shadowed-violet indicates the hydrophobic “hot-spot”). The predicted binding poses of proline (red), CV (violet), loratadine (blue), cyproheptadine (yellow), olanzapine (dark-grey), clofazimine (green) and dapsone (pink) are shown in the figure. (b) Detail of the residues predicted to interact with CV (right side) and proline (left side).

**Table 1.** Compounds obtained by similarity screening and molecular docking analysis. All the drugs obtained by similarity-based screening, in addition to the reference compound (CV) and the natural substrate of TcAAAP069 (PRO), are listed in the table. Columns indicate the compound abbreviation (Abbr.), the popular name and one of the brand names obtained from the website Drugs.com (https://www.drugs.com/), the ZINC database ID, the lowest binding energy (ΔG all), and the lowest binding energy of the most populated cluster (ΔG MPC). Binding energy values were calculated using the AutoDock software. Dash (-) instead of brand name indicates that the compound is marketed under its chemical (popular) name.

| Abbr | Compound (popular name) | Compound (brand name) | ZINC ID | $\Delta G$ all (kcal.mol<sup>-1</sup>) | $\Delta G$ MPC (kcal.mol<sup>-1</sup>) |
| --- | --- | --- | --- | --- | --- |
| CFZ | Clofazimine | Lamprene® | 17953024 | -9.60 | -9.60 |
| LTD | Loratadine | Alavert® | 537931 | -8.01 | -7.25 |
| CPH | Cyproheptadine | Periactin® | 968264 | -7.80 | -7.76 |
| OLZ | Olanzapine | ZyPREXA® | 52957434 | -5.81 | -5.34 |
| CV | Crystal Violet | - | 13763987 | -5.26 | -4.54 |
| **PRO** | **Proline** | - | **895360** | **-4.07** | **-3.60** |
| DPS | Dapsone | Aczone® | 6310 | -1.35 | -1.08 |

The PRO site refers to the groove occupied by the proline pose, comprised by the sequential group of residues 82-87 (GNVGSA), and the CV site refers to the pocket occupied by the CV pose that includes the residues 264-265 (MM), 397-404 (SLIGALPA), 407 (M), and 462-474 (IMVVGTLSMVKGL) (Fig 3b). All the docked poses of CV analogues are located within the CV site (S2 Fig). There is a “hot-spot” rich in A, M and V residues in the CV pocket, where the hydrophobic rings of LTD, CPH, CFZ, and CV appear to converge. OLZ is not near this hot-spot and it binds to the V and A residues of the PRO site, as well as to the residue I-399 of the CV site. As mentioned, DPS predicted poses did not occupy any site inside the protein channel. All these results explain the most probable interaction between the CV, CV analogues and TcAAAP069; however they were not taken as a criterion for the selection of the compounds to be used in the subsequent *in vitro* tests.

### Inhibition of proline transport by CV chemical analogues

The four selected drugs that were structurally similar to CV and the negative control (DPS) were tested as TcAAAP069 inhibitors. Preliminarily, proline transport assays were performed using drug concentrations of 25 μM and 100 μM. All the CV analogues produced a significant proline transport inhibition at both concentrations in epimastigotes, except for OLZ which only inhibited proline transport at 100 μM (S3 Fig). DPS did not inhibit the proline transport in any concentration assayed. Next, the concentration required to inhibit 50% of the proline transport activity (IC_50_) was calculated for the four CV structural analogues. The IC_50_s obtained were 7.1 μM ± 0.8, 23.2 μM ± 3.8, 71.9 μM ± 11.2 and 4.3 μM ± 0.8 for CV, LTD, CPH, and CFZ, respectively (Fig 4). It was not possible to calculate an IC_50_ for OLZ since this compound produced the death of parasites in the lapse time of the assay in concentrations over 300 μM.

**Fig 4.**
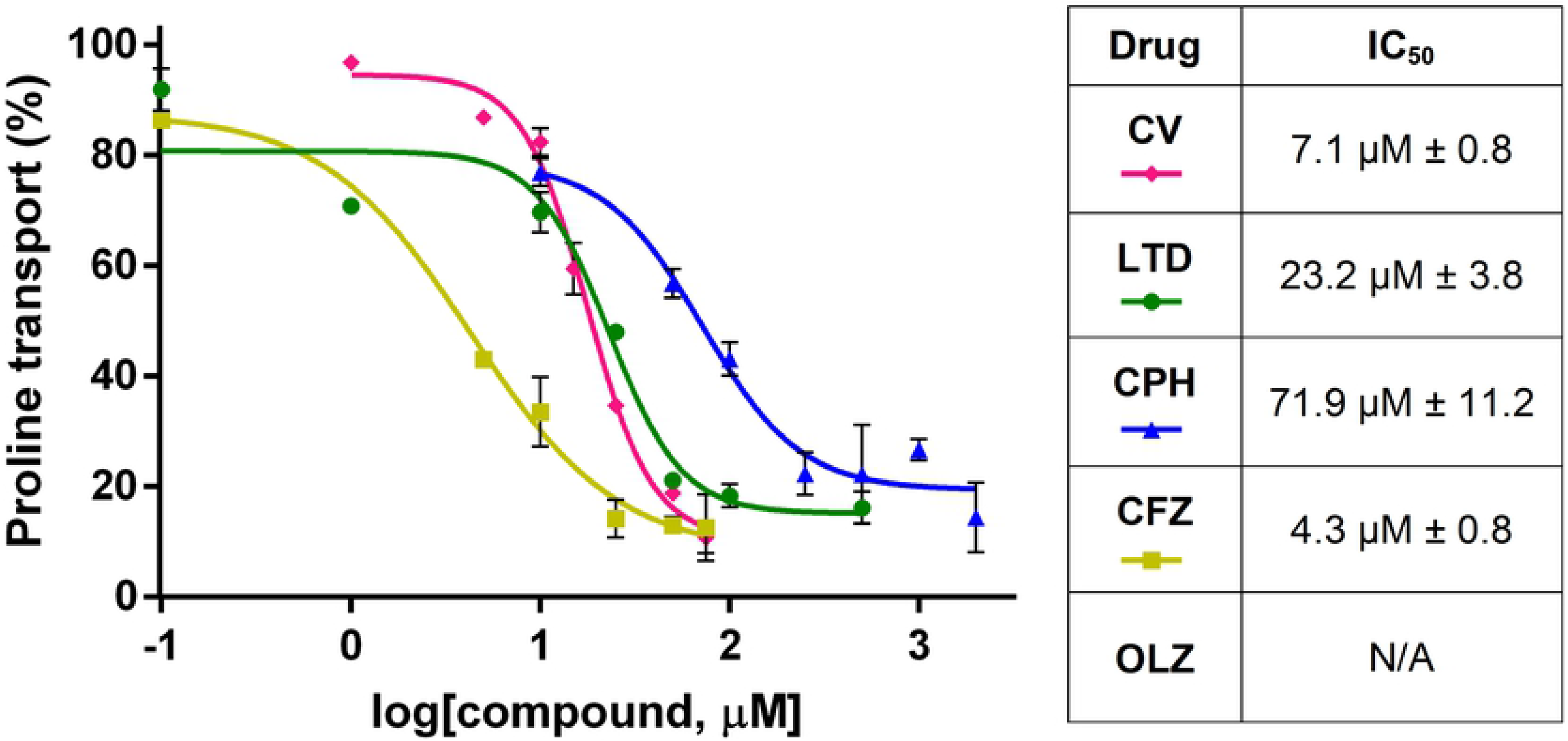
Calculation of CV structural analogues concentrations required to inhibit 50% of proline transport activity (IC_50_). The IC_50_s were calculated for three of the four CV analogues. It was not possible to calculate an IC_50_ for OLZ since this compound produced the death of parasites in the lapse time of the assay in concentrations over 300 μM. The data is expressed as the mean ± standard deviation and corresponds to one of three independent experiments. CV, crystal violet. LTD, loratadine. CPH, cyproheptadine. OLZ, olanzapine. CFZ, clofazimine. N/A, not available.

### Trypanocidal effect of CV structural analogues

Since the four CV analogues tested showed a significant inhibition of proline transport through TcAAAP069 permease, all of them were also evaluated for trypanocidal activity on *T. cruzi* epimastigotes. Although the IC_50_ of OLZ for proline transport inhibition could not be calculated, it was included it in the trypanocidal activity assays because it produced the death of parasites in the proline transport inhibition assay. Parasites were incubated with different concentration of each compound during 48 h and the concentration required to inhibit 50% of parasite growth (IC_50_) was calculated. All the compounds showed trypanocidal activity on epimastigotes, being CFZ the compound with the lowest IC_50_ (9.3 μM ± 0.6) (Table 2 and S4 Fig). Moreover, CFZ presented a significantly lower IC_50_ than the control drug benznidazole (BZL, p<0.001) while the other CV analogues, LTD, CPH and OLZ, were less effective than BZL (p<0.0001 for all of them). Subsequently, the drugs were tested on trypomastigotes, one of the mammalian stages of *T. cruzi*, and all of them proved to have trypanocidal action. CFZ was also the compound that presented the highest effect (2.8 μM ± 0.1) and, together with CPH, showed an IC_50_ value significantly lower than BZL (p<0.0001 and p<0.01, respectively) (Table 2 and S4 Fig). On the contrary, LTD presented a similar IC_50_ value than BZL while OLZ was less effective (p<0.0001). In concordance with the proline transport inhibition assays, OLZ was discarded for further studies due to its elevated IC_50_s (IC_50_ > 100 μM in epimastigotes and IC_50_ > 50 μM in trypomastigotes). The remaining CV analogues were also evaluated on amastigotes, the intracellular mammalian stage of *T. cruzi*, and all of them presented trypanocidal effect with similar (LTD and CPH) or significantly lower IC_50_ values than BZL (CFZ, 1.1 μM ± 0.1, p<0.0001) (Table 2 and S4 Fig). Finally the toxicity of LTD, CPH and CFZ was assessed in VERO cells in order to calculate the selectivity index (SI) and all the CV analogues assayed were more selective to parasite cells with SI values between 5.5 and 36.1 (Table 2).

**Table 2.** Trypanocidal effect of CV chemical analogues in *T. cruzi* Y strain.

| Compound | Epimastigotes | Trypomastigotes | Amastigotes | Selectivity Index |
| --- | --- | --- | --- | --- |
| Benznidazole | $20.9 \mu\text{M} \pm 1.4$ | $14.9 \mu\text{M} \pm 1.5$ | $11.4 \mu\text{M} \pm 0.5$ | - |
| Crystal Violet | $12.7 \mu\text{M} \pm 2.1$ | $0.4 \mu\text{M} \pm 0.0$ | $0.3 \mu\text{M} \pm 0.1$ | - |
| Loratadine | $26.4 \mu\text{M} \pm 1.0$ | $12.9 \mu\text{M} \pm 0.8$ | $13.2 \mu\text{M} \pm 2.4$ | 5.5 |
| Cyproheptadine | $52.6 \mu\text{M} \pm 2.9$ | $11.3 \mu\text{M} \pm 0.6$ | $10.7 \mu\text{M} \pm 1.3$ | 9.1 |
| Olanzapine | $120.5 \mu\text{M} \pm 11.4$ | $54.1 \mu\text{M} \pm 2.9$ | N/A | N/A |
| Clofazimine | $9.3 \mu\text{M} \pm 0.6$ | $2.8 \mu\text{M} \pm 0.09$ | $1.1 \mu\text{M} \pm 0.1$ | 36.1 |

N/A: Data not available

Altogether, the results indicate that CFZ is the only CV analogue tested that presents lower IC_50_ values than BZL in all the *T. cruzi* stages assayed, while LTD and CPH have trypanocidal activity against amastigotes and trypomastigotes with at least similar effect to BZL.

### Trypanocidal effect of CV analogues on other strains

The results presented so far were obtained from *T. cruzi* parasites from Y strain (discrete typing unit –DTU– TcII). The trypanocidal action of LTD, CPH and CFZ was also evaluated on epimastigotes of two other DTU strains, such as Dm28c (DTU TcI) and CL Brener (DTU TcVI) since variations in trypanocidal treatment response have been reported for different *T. cruzi* strains (45). BZL and CV were assayed as well in order to compare the trypanocidal effect between strains. The Y parasites were 1.6- and 1.3-fold more sensitive to BZL treatment than Dm28c and CL Brener epimastigotes, respectively (p<0.0001, Table 3). Dm28c was 1.2-fold more resistant to BZL than CL Brener (p<0.01). On the contrary, Dm28c and CL Brener parasites presented similar response to CV treatment and were 31.8- and 4.0-fold, respectively, more sensitive to this compound than Y epimastigotes (p<0.0001). In addition, the results indicate that the three CV chemical analogues are effective in other *T. cruzi* strains, including DTUs of clinical relevance (46) (Table 3 and S5 Fig). Dm28c parasites were significantly more sensitive to LTD and CPH than the epimastigotes of Y (p<0.01 and p<0.0001, respectively) and CL Brener strains (p<0.0001, for both drugs). However, no differences between strains were observed in the treatment with CFZ.

**Table 3.** Trypanocidal effect of CV structural analogues in epimastigotes from different strains of *T. cruzi*.

| Compound | *T. cruzi* strain |  |  |
| --- | --- | --- | --- |
|  | Y | Dm28c | CL Brener |
| Benznidazole | 20.9 $\mu\text{M} \pm 1.4$ | 32.5 $\mu\text{M} \pm 1.9$ | 27.8 $\mu\text{M} \pm 1.8$ |
| Crystal Violet | 12.7 $\mu\text{M} \pm 2.1$ | 0.4 $\mu\text{M} \pm 0.0$ | 3.2 $\mu\text{M} \pm 1.1$ |
| Loratadine | 26.4 $\mu\text{M} \pm 1.0$ | 21.0 $\mu\text{M} \pm 1.2$ | 28.6 $\mu\text{M} \pm 2.0$ |
| Cyproheptadine | 52.6 $\mu\text{M} \pm 2.9$ | 37.6 $\mu\text{M} \pm 0.7$ | 50.6 $\mu\text{M} \pm 2.2$ |
| Olanzapine | 120.5 $\mu\text{M} \pm 11.4$ | N/A | N/A |
| Clofazimine | 9.3 $\mu\text{M} \pm 0.6$ | 7.5 $\mu\text{M} \pm 0.8$ | 9.0 $\mu\text{M} \pm 0.5$ |

N/A: Data not available

### Synergism between CV structural analogues and benznidazole

The synergistic action between BZL, LTD, CPH, CFZ and a combination of these three CV analogues (LTD-CPH-CFZ) was evaluated in *T. cruzi*. Drug interactions were classified according to the combination index (CI) recognized as the standard measure of combination effect based on the Chou-Talalay method (44). CI < 1 means synergism, CI = 1 indicates additive effect, and CI > 1 is interpreted as antagonism. The IC_50_ calculated for BZL treatment alone was 18.57 μM ± 1.07 and for the LTD-CPH-CFZ treatment was 18.05 μM ± 0.96. While each compound by itself did not enhance the BZL trypanocidal activity (data not shown), the combined administration of LTD-CPH-CFZ and BZL produced a synergistic effect. All combinations between BZL and LTD-CPH-CFZ presented a CI < 1 with different synergism degree, from strong to slight synergism (Fig 4a). The exceptions were the combination of 1/2 IC_50_ LTD-CPH-CFZ with 0.1 and 1 μM BZL, which presented a nearly additive effect and moderate antagonism, respectively. The Chou-Talalay plot shows the CI and the effect (Fa, fraction affected) produced by each combination point, where 0 means no effect and 1.0 means maximum trypanocidal effect (Fig 4b). Dose-reduction index (DRI) represents the dose reduction that is allowed in combination for a given degree of effect as compared with the dose of each drug alone. DRI > 1 indicates favorable dose-reduction. Except for the two combinations that did not present a synergistic effect, the DRI values were higher than 1 for all the combinations assayed with values between 2.5 and 374.2 for BZL and between 1.2 and 10.4 for LTD-CPH-CFZ (S1 Table). Altogether, these results suggest that combination therapy improves the trypanocidal action of the CV analogues together with BZL.

**Fig 5.**
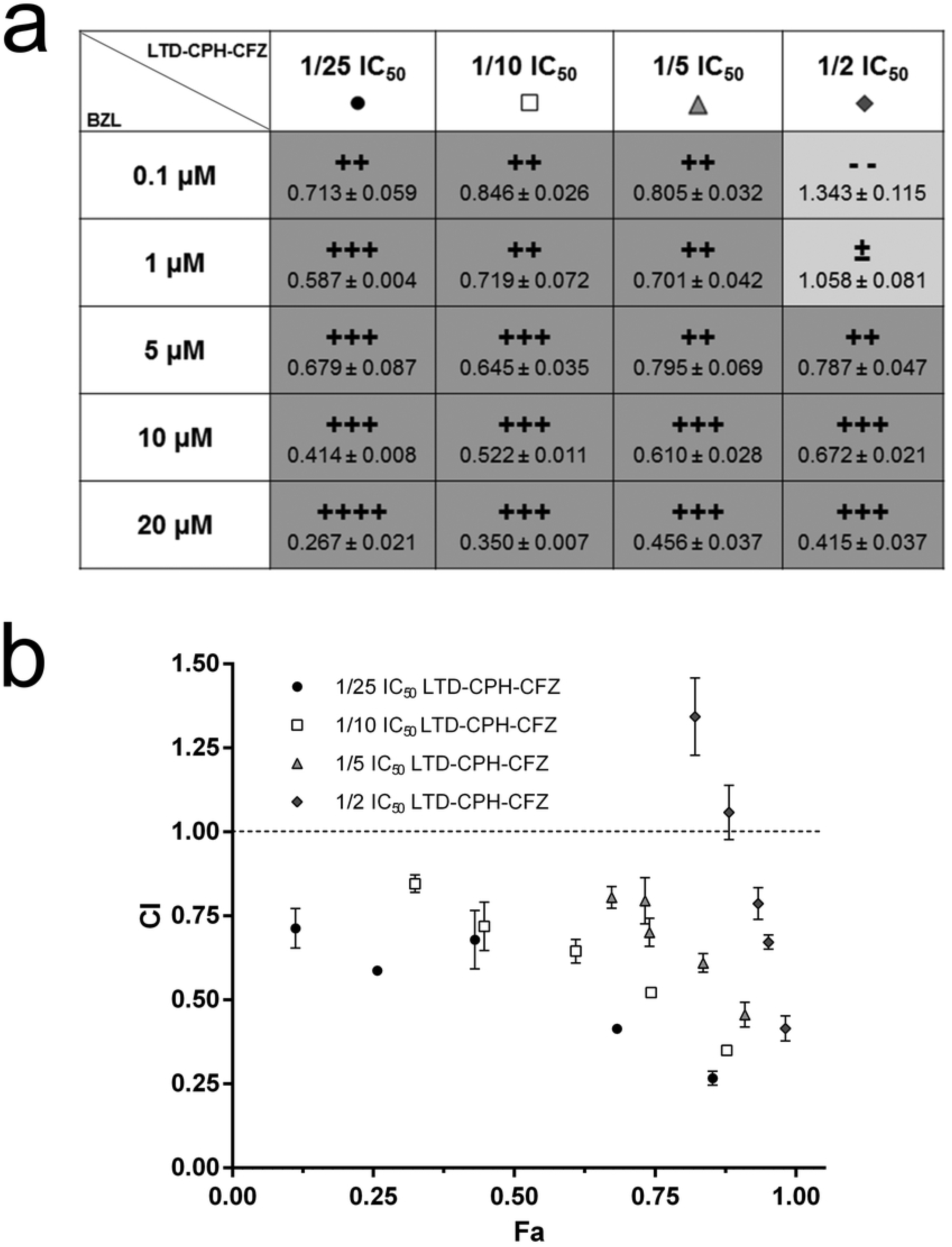
Synergism between benznidazole and the combination of the crystal violet analogues in *T. cruzi* epimastigotes. (a) Drug combinations between BZL and CV analogues (LTD-CPH-CFZ). Combination index (CI) value for each combination point is presented under the corresponding graded symbol. Graded symbols mean strong synergism (++++, CI between 0.1-0.3), synergism (+++, CI between 0.3-0.7), moderate synergism (++, CI between 0.7-0.85), nearly additive effect (±, CI between 0.9-1.1), and moderate antagonism (− −, CI between 1.2-1.45) (47). The boxes coloured with light-grey correspond to the combination points where no synergism was observed. (b) Chou-Talalay plot. Representation of CI vs effect (Fa, fraction affected), where CI > 1, CI = 1 (dotted line) and CI < 1 indicate antagonism, additive effect and synergism, respectively. For each data series BZL concentrations increase from left to right (from 0.1 to 20 μM). The data is expressed as the mean ± standard deviation and corresponds to one of three independent experiments. All calculations were performed with CompuSyn software. LTD, loratadine. CPH, cyproheptadine. CFZ, clofazimine. BZL, benznidazole. IC50 LTD = 25 μM. IC50 CPH = 50 μM. IC50 CFZ = 10 μM. LTD-CPH-CFZ, combination of the three crystal violet analogues as a single drug. 1/2 IC50, refers to the sum of half of each IC50, 12.5 μM + 25 μM + 5 μM = 42.5 μM. 1/5 IC50, 5 μM + 10 μM + 2 μM = 17 μM. 1/10 IC50, 2.5 μM + 5 μM + 1 μM = 8.5 μM. 1/25 IC50, 1 μM + 2 μM + 0.4 μM = 3.4 μM.

## Discussion

New alternatives to current treatments against Chagas disease have been explored in the last decade, including new trypanocidal drugs, such as the posaconazole, and the use of BZL in the Chagas cardiomyopathy (BENEFIT trial) (48–51). Unfortunately none of them were successful, and there is still an urgent need for the development of new therapeutic substitutes for conventional treatments. Drug repurposing or repositioning consist of finding new indications for drugs already approved for its use in humans. This approach is a recommended strategy by the World Health Organization (WHO) to fight neglected diseases, like Chagas disease, since costs and development time are significantly reduced for its application in therapy.

There are some recent examples of drug repositioning by virtual screening that use the *T. cruzi* polyamine transporter TcPAT12 as main target; triclabendazole, an antiparasitic, sertaconazole, a broad-spectrum antifungal, paroxetine, an antidepressant agent, isotretinoin, a well-known medication used to treat severe acne, and cisapride, a gastroprokinetic agent. All these compounds proved to be effective as TcPAT12 inhibitors that also have trypanocidal action (52–54). Regarding proline transport as therapeutic target, to our knowledge there are no previous reports of drug repositioning although some proline analogues have been studied as trypanocidal compounds. For example, the proline analogue L-thiazolidine-4-carboxylic acid (T4C) has been reported as a proline transporter inhibitor with trypanocidal effect (55), and more recently, another proline analogue synthetized by rational design and called ITP-1G has been validated as a TcAAAP069 inhibitor that also possesses trypanocidal activity (25).

Other compounds have been identified by drug repositioning as possible trypanocidal drugs using different targets; such as the cruzipain (56–59), some *T. cruzi* kinases (60–62) and the trans-sialidase (63), among other examples. Other drug repurposing approaches have inferred the possible trypanocidal activity of already-approved drugs based on the reported effect or mechanism of action of some compounds in other organisms (64–69).

In this work, a combined strategy involving similarity-based virtual screening followed by *in vitro* assays was applied in order to obtain new trypanocidal compounds from a database that contains already-approved drugs. CV was selected as a reference molecule since it has a reported trypanocidal action and one of its mechanisms of action involved proline transport inhibition (4). The aim was to obtain structurally related compounds which may have similar biological activity but with no (or less) toxicity toward the host cells since they are already in use to treat human diseases. The proline transporter TcAAAP069 was selected as the probable main target because: a) proline is a relevant amino acid for *T. cruzi* biology, including host cell invasion and life cycle progression (16,17), b) TcAAAP069 mediates proline uptake and it can also be used as a way of entry drugs into the parasite and, c) there is evidence of compounds that inhibit TcAAAP069 and also have trypanocidal activity, like the proline analogues T4C and ITP-1G (25,55).

Once CV was validated as a TcAAAP069 inhibitor, the similarity-based virtual screening was performed and four drugs already-approved for clinical use were obtained for further analysis. All these CV chemical analogues retrieved by the virtual screening have already been tested as antiparisitic agents in protozoans. LTD was evaluated in *Leishmania* (L.) *infantum* as an antileishmanial compound and even though it presented an intermediate effect on promastigotes it did not have effect on intracellular amastigotes (70). CPH was assayed in *T. evansi* in order to evaluate the ability of this compound to reverse drug resistance to some trypanocidal drugs, like suramin, and the initial experiments showed the intrinsic toxic effect of this compound (71). OLZ was tested in *Toxoplasma gondii* due to the strong link demonstrated between toxoplasmosis and psychiatric disorders but it presented a low anti-toxoplasmic activity (72). Finally, CFZ was recently identified as a cruzipain inhibitor in *T. cruzi* with trypanocidal action, but it was also reported that cruzipain is not the only target for this compound (73).

LTD, CPH and CFZ were all able to inhibit proline transport and also proved to have trypanocidal effect in epimastigotes, trypomastigotes and amastigotes of the parasite *T. cruzi*. The IC_50_s obtained for these CV analogues were in the range of those reported with BZL in epimastigotes for strains from different DTUs (IC_50_ values between 7-30 μM) (45). Moreover, the trypanocidal effect of CFZ was significantly higher than the observed with BZL in trypomastigotes and amastigotes while the other two CV analogues showed at least similar effect to the BZL. In contrast, OLZ presented an elevated IC_50_ both for epimastigotes and trypomastigotes, and it was discarded for further analysis. The molecular docking predicted two hydrophobic sites where CV and proline could stably bind to the transporter TcAAAP069 and, interestingly, all the CV analogues clustered inside the TcAAAP069 channel, mainly in the CV site as expected for structural analogues. These results could explain the observed inhibition of proline transport by the CV structural analogues.

The fact that LTD, CPH and CFZ were also effective as trypanocidal compounds on epimastigotes of CL Brener and Dm28c strains is interesting because not all *T. cruzi* strains present the same response to current treatments (45), and even drug resistance has been strain-associated (74). Indeed there were differences on the treatment response to BZL and CV between the three strains evaluated in this work. The response observed to CV was particularly striking, with IC_50_ values 4.0- and 31.8-fold lower for CL Brener and Dm28c epimastigotes, respectively, than the IC_50_ obtained for the Y strain. These results indicate the importance of evaluate trypanocidal compounds in several strains in order to prevent different and unexpected responses. Regarding the CV analogues, CFZ presented the same effect on the three strains evaluated, while LTD and CPH proved to be significantly more effective on Dm28c strain.

The combination of two or more drugs with different mechanisms of action is an alternative approach to increase the success rate of drug repositioning and, although multidrug therapy may be difficult to implement, combination of drugs has been used to treat other neglected diseases, for example the leprosy multidrug treatment recommended and supplied by the WHO involves rifampicin, clofazimine and dapsone (MDT-COMBI®). The combination of already-approved drugs enables dosage reduction and diminishes toxicity, in addition to an improved efficacy and reduced drug resistance emergence. The final experiment herein presented showed positive interactions between the well-known drugs LTD, CPH and CFZ, and BZL in epimastigotes of *T. cruzi*. Most of the combination points of BZL with LTD-CPH-CFZ presented a synergistic effect, with CI values from 0.27 to 0.85 and DRI values from 2.5 to 374.2 for BZL and between 1.2 and 10.4 for LTD-CPH-CFZ. For example, the combination of 1 μM BZL with 1/5 IC_50_ LTD-CPH-CFZ (17 μM, CI = 0.70) produced an effect of 74% with 54-fold less BZL and 1.5-fold less LTD-CPH-CFZ than each drug alone needed to achieve the same effect. These results suggest that the combination therapy would make possible to diminish the dosage of BZL and thus, the toxicity and side effects reported (6). However, future studies will need to focus on the effect of the combination therapy between BZL and the CV structural analogues in trypomastigotes and in infected cells.

The final goal would be to identify new treatments based on drug repositioning and/or combination therapy that are more effective, better tolerated, and simpler to administrate than current therapies for treating Chagas disease. Taking into account the urgent need for the development of new drugs for Chagas disease, we can conclude that CV chemical analogues could be a starting point to design therapeutic alternatives either alone or in combination with BZL to treat this neglected disease.

## Acknowledgments

Special thanks to Lic. Fabio di Girolamo (IDIM, UBA-CONICET), Dr. Patricia Bustos and Dr. Alina Perrone (Instituto Nacional de Parasitología “Dr. Mario Fatala Chaben”) for technical support, and to Dr. Viviana Amparo Ybarra and Dr. Guillermo Buiyi Alonso (INGEBI-CONICET) for their collaboration. Also we would like to thank Thomas Lemker from “Inte:Ligand Software Development & Consulting” for his help with the license of the LigandScout and to Richmond Laboratories for olanzapine supply. CAP and MRM are members of the career of scientific investigator; MS, EVV and CR are research fellows from CONICET, LG is a undergraduate student fellow

## Supporting Information

**S1 Fig. Chemical features shared between crystal violet and the selected compounds.**

The LigandScout software was used to identify the common chemical features between crystal violet (CV) and its chemical analogues. This algorithm performs feature-based structure alignments where the similarities are calculated as the number of matched feature pairs (MFP). (a) In order to perform the structure comparisons, the 8 CV features were set as references. (b) The LigandScout similarity score and the number of MFP obtained for each compound are listed. (c) The features shared between CV and each compound are shown and the structure alignments with the van der Waals surfaces are schematized. N/A, not available. AR, aromatic ring (purple circles). H, hydrophobic area (yellow remarks). PI, positive ionizable atom (purple lines).

**S2 Fig. Predicted poses by molecular docking of the crystal violet structural analogues and the proline permease TcAAAP069.**

Residues corresponding to the PRO and CV sites in TcAAAP069 are indicated in green and violet, respectively. Detail of the TcAAAP069 residues predicted to interact with (a) clofazimine, (b) loratadine, (c) cyproheptadine and (d) olanzapine.

**S3 Fig. Inhibition of proline transport by crystal violet chemical analogues in wild type parasites (TcWT).**

The crystal violet analogues and dapsone were evaluated as potential proline transport inhibitors at two concentrations, 25 and 100 μM. Control, no treatment. The data is expressed as the mean ± standard deviation and corresponds to one of three independent experiments. DPS, dapsone. LTD, loratadine. CPH, cyproheptadine. OLZ, olanzapine. CFZ, clofazimine. *, p< 0.05; ***, p< 0.001; ****, p<0.0001. ns, not significant.

**S4 Fig. Trypanocidal effect of CV structural analogues in (a) epimastigotes, (b) trypomastigotes and (c) amastigotes of***T. cruzi* **Y strain.**

The concentrations required to inhibit 50% of parasite growth or parasite survival were calculated for the four CV chemical analogues. The data is expressed as the mean ± standard deviation and corresponds to one of three independent experiments. BZL, benznidazole. CV, crystal violet. LTD, loratadine. CPH, cyproheptadine. OLZ, olanzapine. CFZ, clofazimine. N/A, not available.

**S5 Fig. Trypanocidal effect of CV structural analogues concentrations in epimastigotes of (a)***T. cruzi* **DM28c and (b) CL Brener strains.**

The concentrations required to inhibit 50% of parasite growth were calculated for three CV chemical analogues. OLZ was not tested in these strains because of the high IC_50_s values obtained in trypomastigotes and epimastigotes of the Y strain. The data is expressed as the mean ± standard deviation and corresponds to one of three independent experiments. BZL, benznidazole. CV, crystal violet. LTD, loratadine. CPH, cyproheptadine. OLZ, olanzapine. CFZ, clofazimine. N/A, not available.

**S1 Table. Dose-reduction index for drug combination therapy with benznidazole and crystal violet chemical analogues in wild type epimastigotes of T. cruzi Y strain.**

